# Peptide-level assay of protein interaction using combinatorial native and denatured proteolysis

**DOI:** 10.64898/2026.11.09.765259

**Authors:** Siyuan Chen, Zhixin Tian

## Abstract

Protein–protein interactions underlie many extracellular defense and protease-regulatory processes, yet identifying their structural interfaces directly in complex biological samples remains challenging. Limited proteolysis coupled with mass spectrometry provides peptide-level readouts of local protease accessibility, but linking condition-dependent peptide recovery to structural regions in endogenous proteomes requires complementary analysis. Here, we report peptide-level assay of protein interaction (PLAPI) using combinatorial native and denatured proteolysis. A single human serum specimen was analyzed in three paired technical preparations using brief trypsin digestion under native conditions and extended digestion after denaturation, followed by isotopic diethyl labeling and LC-MS/MS. Applying a 2-of-3 detection rule yielded 920 denatured-only peptides and 124 peptides detected under both conditions, with no native-only peptides. Mapping to the reference structures placed 13 denatured-only C3b peptides at the Factor B-binding interface and identified multiple denatured-only A2M intersubunit contact regions. Within the A2M bait region, adjacent peptides exhibited different detection patterns; two were detected under both conditions, whereas one was detected only after denaturation. STRING association and enrichment analyses linked the candidate proteins to extracellular defense and protease regulation. These results indicate that PLAPI can provide peptide-level evidence for prioritizing structural regions in complex serum samples and guide focused follow-up analyses.

## INTRODUCTION

Protein–protein interactions (PPIs) assemble proteins into complexes and coordinate their activities in processes such as signal transduction, immune responses, metabolism, and protein homeostasis^1,2^. According to human interactome studies, disruptions in PPI networks can impair protein complex function and contribute to disease^3–5^. Understanding PPIs is therefore important for elucidating the molecular mechanisms of disease^3,6^. At the molecular level, protein–protein binding interfaces mediate partner recognition and complex assembly. Their structural and chemical features help determine interaction specificity and stability^7^. Protein–protein binding interfaces thus offer an important entry point for understanding how altered PPIs contribute to disease and identifying potential sites for therapeutic intervention^4,5,8^. However, mapping protein–protein binding interfaces directly in complex biological samples remains challenging^6,9^.

Limited proteolysis offers a way to examine protein surfaces within complex samples by measuring local protection from cleavage. When part of a protein surface is shielded, protease access is restricted and cleavage is reduced; whereas exposed, flexible segments are more readily cleaved^10^. Coupling limited proteolysis to mass spectrometry allows these differences in cleavage and peptide release to be resolved at the peptide level. This principle has been widely applied to protein-ligand interaction studies. Lomenick et al. established drug affinity responsive target stability (DARTS), using ligand-induced resistance to proteolysis to identify small-molecule target proteins^11^. Feng et al. developed limited proteolysis-coupled mass spectrometry (LiP-MS), enabling peptide-level measurements of local structural changes in complex proteomes^12,13^. Li et al. subsequently introduced the peptide-centric local stability assay (PELSA), using rapid digestion of native proteins to detect ligand-induced changes in local protein stability^14^.Because bound ligands and protein partners can both shield local regions and stabilize their conformations, these studies provide a mechanistic basis for extending proteolytic accessibility measurements to PPI interface analysis.

Existing PPI studies have used limited proteolysis to locate binding-sensitive regions by comparing proteins in different binding states. Dörig et al. developed FLiP-MS to compare complex-bound and monomeric protein populations by combining serial ultrafiltration with LiP-MS. The resulting marker library was enriched in peptides mapping to protein-binding interfaces^15^.Holfeld et al. compared cell extracts exposed to different concentrations of a purified bait protein, identifying candidate interactors and localizing known binding regions at peptide resolution^16^.A complementary strategy is therefore needed to identify regions with restricted peptide release under native conditions directly in complex samples and prioritize them for interface mapping, without requiring prior separation of assembly states or the addition of purified bait proteins.

To address this need, this study established the PLAPI workflow for human serum, pairing brief digestion at a high enzyme-to-substrate ratio under native conditions (Native) with extended digestion after denaturation (Denatured). The aim was to determine whether this comparison could provide reproducible peptide-level information relevant to protein interfaces and functional regions. Peptide detection and quantitative profiles were compared across paired replicates. The resulting detected patterns were then examined in reference structures of C3b–Factor B and A2M, which provide defined assembly interfaces and a protease-sensitive bait region for interpretation. These analyses linked Denatured-only peptides to subunit contacts and identified different detection patterns among neighboring bait-region segments, demonstrating the application of the workflow to local structural analysis in serum.

## EXPERIMENTAL SECTION

### Serum Samples and Experimental Design

A single human serum specimen was used to prepare three paired Native– Denatured technical replicates. Each preparation contained 2 μL of serum. The serum protein concentration was 63 mg/mL, as measured by a BCA assay, corresponding to approximately 126 μg of protein per aliquot. Human serum samples were obtained from the same cohort of HBV-related hepatocellular carcinoma patients as described in our previous work ^17^. The study was approved by the Ethics Committee of Zhongshan Hospital (approval no. B2019-100R) and conducted in accordance with the Declaration of Helsinki.

### Native-State Limited Proteolysis

For each Native preparation, 2 μL of serum was diluted with 98 μL of PBS. Trypsin was added at an enzyme-to-substrate ratio (E:S) of 1:2 (w/w). Samples were vortex-mixed for 10 s and incubated at 37 °C for 50 s; the incubation time began after mixing. Proteolysis was quenched by adding 489 μL of 8 M guanidine hydrochloride (GdmCl) in 60 mM HEPES, pH 8.2, to give a final GdmCl concentration of 6 M^14^. DTT was then added to 10 mM, and the sample was incubated at 55 °C for 30 min. After cooling to room temperature, IAA was added to 20 mM, followed by 30 min of incubation in the dark. An equal amount of DTT was added, and the sample was incubated for a further 30 min at room temperature.

### Denatured-State Digestion

For each paired Denatured preparation, 8 μL of 8 M urea was added to 2 μL of serum. Samples were vortex-mixed for 10 s and incubated at room temperature for 1 h. The 10 μL mixture was brought to 100 μL with 90 μL of PBS, resulting in a 50-fold dilution of serum and a final urea concentration of 0.64 M. Samples were reduced with 10 mM DTT at 55 °C for 30 min, alkylated with 20 mM IAA for 30 min in the dark at room temperature, and treated with an equal amount of DTT for 30 min at room temperature. Trypsin was added at an E:S ratio of 1:50 (w/w), and digestion proceeded at 37 °C for 16 h. Digestion was terminated by adding TFA to a final concentration of 0.1% (v/v), followed by vortex mixing for 10 s.

Both Native and Denatured preparations were processed using 10 kDa centrifugal filter units at 14,000 × g for 1 h. The membrane was washed with 200 μL of 60 mM HEPES, pH 8.2, and centrifuged again at 14,000 × g for 1 h^14^. The filtrates were combined and acidified by adding TFA at 1% of the filtrate volume before C18 solid-phase extraction.

### Isotope-Coded Diethyl Labeling

Desalted Denatured peptides were labeled with light acetaldehyde (CH₃CHO), and Native peptides were labeled with heavy [U– ¹³C₂]acetaldehyde (¹³CH₃¹³CHO) by reductive diethylation. The labels were introduced at peptide N termini and lysine residues. The net mass additions were +56.0626 Da for light labeling and +60.0760 Da for heavy labeling per modified amino group. Corresponding light– and heavy-labeled preparations were directly combined at 1:1 (w/w) ratio before LC–MS/MS analysis^18^.

### LC–MS/MS and Database Search

Paired peptide mixtures (5 μg per analysis) were analyzed by LC–DDA– MS/MS using a Dionex Ultimate 3000 RSLCnano system coupled to an nano-ESI Q Exactive mass spectrometer. Peptides were loaded onto a self-packed C18 trap column (5 cm × 200 μm i.d., 360 μm o.d.) packed with Phenomenex Jupiter C18 particles (5 μm, 300 Å) at 5 μL/min for 5 min and separated on a self-packed C18 analytical column (70 cm × 75 μm i.d., 360 μm o.d.) packed with the same material at 300 nL/min. Mobile phase A consisted of 99.8% water and 0.2% formic acid, and mobile phase B consisted of 90% acetonitrile, 9.8% water, and 0.2% formic acid. The programmed gradient was 7–42% B over 95 min, 42–95% B over 5 min, 95% B for 3 min, 95–7% B over 2 min, and 7% B for 10 min. The 115-min gradient and re-equilibration sequence, together with the 5-min loading step, constituted a 120-min LC program. The electrospray voltage was 2.8 kV. Raw files were processed using Proteome Discoverer version 1.4.0.288 against a local human plasma protein FASTA database (uniprot_plasma_6274_20201105.fasta). Separate Sequest HT searches were performed for light– and heavy-labeled peptides. Trypsin was specified with up to two missed cleavages. Fixed modifications were light diethylation (+56.063 Da) or heavy diethylation (+60.076 Da) at peptide N termini and lysine residues, and carbamidomethylation (+57.021 Da) at cysteine residues^18^. Peptide lengths of 6–144 residues, a maximum ΔCn of 0.05, and precursor and fragment mass tolerances of 10 ppm and 0.6 Da, respectively, were used. Peptide-spectrum matches were validated using Percolator with strict and relaxed target FDR thresholds of 0.01, based on q-values. Light– and heavy-labeled results were combined after harmonizing label names and removing duplicate peptide sequence/modification entries. Paired light– and heavy-labeled precursor intensities were used to calculate within-preparation Denatured/Native ratios; log₂FC was defined as log₂(Denatured/Native).

### Peptide Classification and Structural Analysis

A peptide was considered detected in a condition when it was observed in at least two of the three preparations. Peptides meeting this criterion in both conditions were classified as Overlap; peptides meeting it only in Denatured were classified as Denatured-only; peptides meeting it only in Native were classified as Native-only. For peptides quantified in both conditions, log₂FC was calculated as log₂(Denatured/Native).

Peptides were mapped to UniProt sequences and reference structures, including C3b–Factor B (PDB 2XWJ) and A2M (PDB 6TAV). A mapped peptide was classified as an interface peptide when at least one peptide non-hydrogen atom was within 4.5 Å of a non-hydrogen atom in a neighboring chain. Structural measurements and calculation definitions are provided in the Supporting Information.

### STRING and Functional Enrichment

Proteins carrying at least one Denatured-only peptide were analyzed using the STRING website for Homo sapiens using a combined-score threshold of ≥0.90. GO and KEGG enrichment results were obtained from the STRING analysis of the 107 mapped proteins. STRING analysis was performed using STRING version 12.5 for Homo sapiens (accessed on November 1, 2026). GO and KEGG enrichment analyses were conducted using the entire genome proteome as the reference background.

## RESULTS AND DISCUSSION

### Comparative Native–Denatured Digestion and Peptide Detection

The PLAPI workflow was established to compare peptide release under native and denaturing conditions and relate these differences to protein structure (**Figure 1**). Aliquots of the same serum sample were processed in three paired replicates, comparing brief digestion at a high enzyme-to-substrate (E:S) ratio under Native conditions (50 s; E:S, 1:2) with extended digestion under Denatured conditions (16 h; E:S, 1:50). Released peptides were recovered by ultrafiltration, differentially isotope-labeled, and combined for liquid chromatography–tandem mass spectrometry (LC–MS/MS). Peptide detection patterns and relative signals were compared between conditions, followed by peptide mapping onto selected protein complex structures.

**Figure 1.**
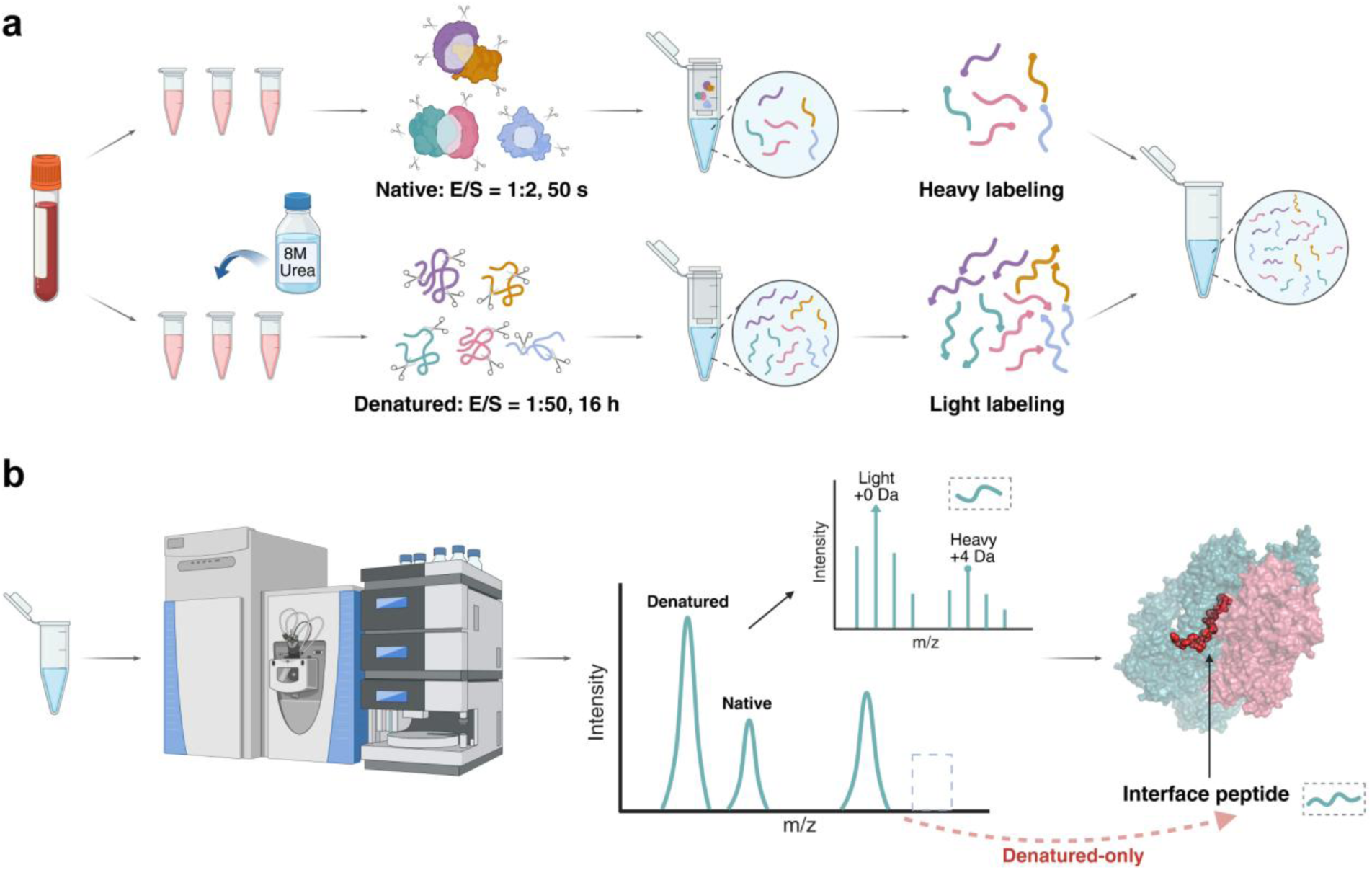
PLAPI workflow and peptide-level structural interpretation. (a) A human serum aliquot is divided into paired Native and Denatured preparations. Native proteins undergo brief trypsin digestion (E:S, 1:2; 50 s), whereas Denatured proteins undergo urea unfolding followed by extended digestion (E:S, 1:50; 16 h). Peptides from both conditions are recovered by ultrafiltration and C18 desalted. Native peptides receive the heavy ¹³C₂-diethyl label, and Denatured peptides receive the light diethyl label; labeled samples are then combined. (b) The mixed peptides are analyzed by LC– MS/MS. The light and heavy precursor ions differ by approximately 4 Da per modified amino group. Peptide detection patterns are compared between conditions and mapped onto reference structures to locate candidate interface-associated peptides. Workflow, spectral, and structural illustrations are schematic; the peak pattern is not a measured mass spectrum.

Across the three paired replicates, peptide counts under Native conditions were 130, 194, and 112, compared with 1,024, 1,113, and 1,056 under Denatured conditions; the corresponding protein counts were 41, 53, and 40 versus 131, 145, and 134, respectively (**Figure 2a, 2b**). Denatured digestion therefore yielded broader peptide and protein coverage in every paired replicate. Quantitative reproducibility was assessed using log₂FC. Pairwise comparisons included peptides quantified under both conditions in both replicates being compared and yielded correlation coefficients of 0.87–0.90 (**Figure 2c–2e**). These correlations indicate that the relative signal differences between the two digestion conditions were consistent across replicates. Together, the coverage and correlation results establish a reproducible contrast between Native and Denatured digestion, providing a basis for classifying peptides by their detection across conditions.

**Figure 2.**
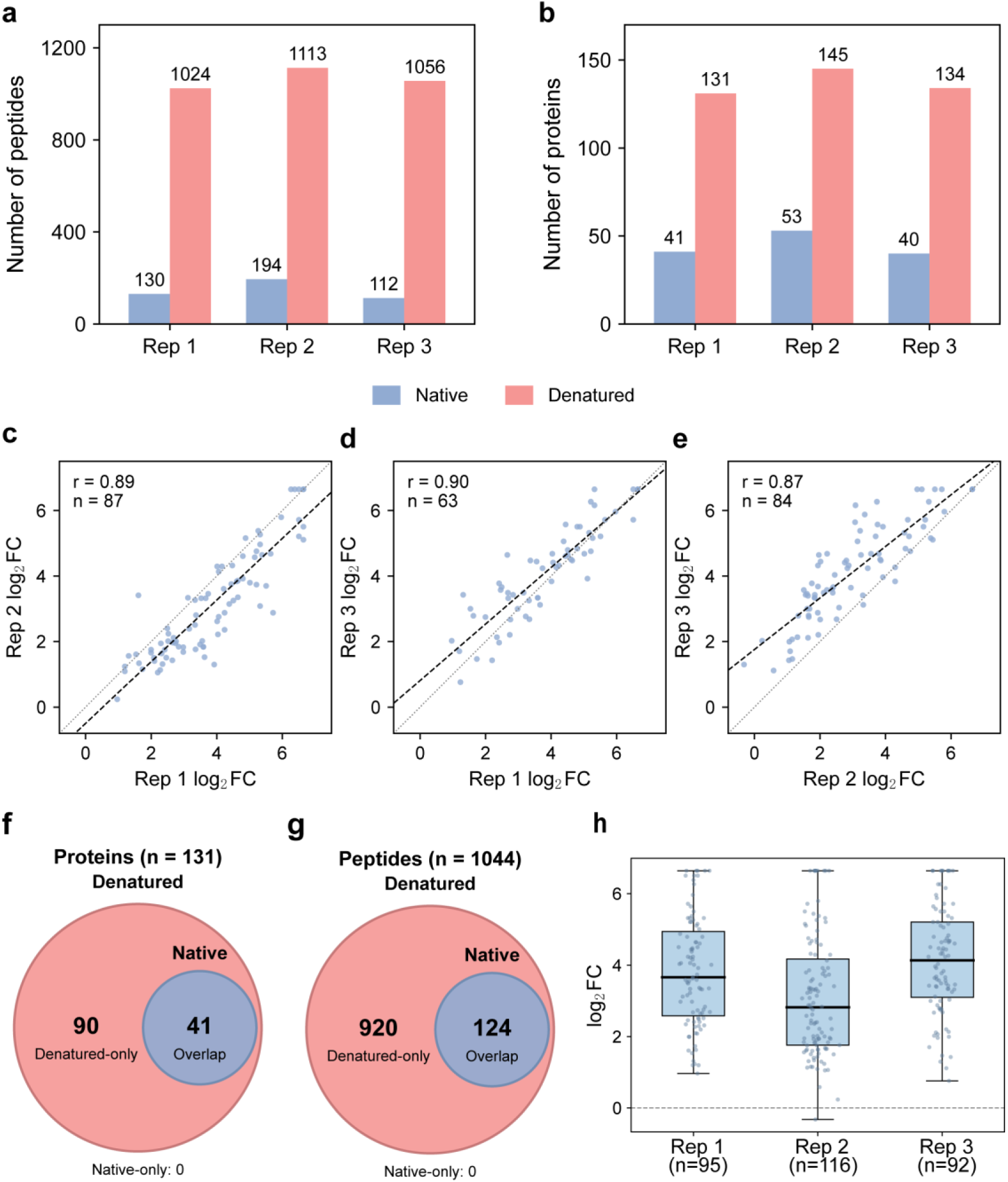
Peptide and protein coverage, quantitative reproducibility, and replicate-based detection classes. (a, b) Numbers of peptides (a) and proteins (b) identified in each Native and Denatured technical preparation. (c–e) Pairwise correlations of log₂(Denatured/Native) values for peptides quantified in both conditions in both preparations being compared: Rep 1 versus Rep 2 (r=0.89, n=87), Rep 1 versus Rep 3 (r=0.90, n=63), and Rep 2 versus Rep 3 (r=0.87, n=84). (f, g) Classification under the 2-of-3 rule for proteins (f) and peptides (g). The protein set contains 90 Denatured-only and 41 Overlap proteins; the peptide set contains 920 Denatured-only and 124 Overlap peptides. No Native-only proteins or peptides met the rule. The circle areas are illustrative and are not proportional to the counts. (h) Paired log₂(Denatured/Native) values for peptides quantified in both channels in each preparation (Rep 1, n=95; Rep 2, n=116; Rep 3, n=92). The dashed horizontal line marks zero. Points represent individual peptide measurements; boxes show the interquartile range and median, and whiskers extend to observations within 1.5 × the interquartile range.

Applying the prespecified 2-of-3 detection rule, yielded 920 Denatured-only peptides (88.1%) and 124 Overlap peptides (11.9%), with no Native-only peptides (**Figure 2f, 2g**; **Supplementary Table S1**). Thus, the reproducibly detected Native peptide set was entirely contained within the Denatured set, while most peptide sequences were reproducibly recovered only under Denatured conditions. Among peptides quantified under both conditions within each paired replicate, log₂FC values were predominantly positive across all three replicates (**Figure 2h**), indicating higher signals under Denatured conditions for most of these peptides. The difference between digestion conditions therefore extended beyond the number of detected sequences to the relative signals of peptides quantified in both conditions. Together, these findings show that brief Native digestion yielded a restricted peptide profile, whereas extended Denatured digestion expanded sequence coverage and increased the measured signals of most peptides in the paired comparisons. This contrast identifies regions with distinct peptide recovery under the two digestion conditions for subsequent analysis in the context of protein assembly interfaces and functional regions.

### Global Peptide Features and C3b–Factor B Interface Mappin

Following peptide classification, global comparisons were performed to assess whether Denatured-only and Overlap peptides differed in their sequence and structural features. Sequence-level summaries, peptide-averaged relative solvent-accessible surface area (mean rSASA)^19^, and normalized Cα B-factors did not consistently distinguish the two classes (**Supplementary Figure S1**; **Supplementary Table S2**). To examine their relationship to local structural environments, both peptide classes were subsequently mapped onto selected protein complexes, focusing on known intersubunit interfaces.

Mapping C3b peptides onto the C3b–Factor B complex identified a Denatured-only subset at regions partially shielded by Factor B. This complex was selected because the C3b dataset contained both detection classes, comprising 64 unique Denatured-only and nine unique Overlap peptide sequences, and PDB 2XWJ^20,21^ provided a resolved assembly with an explicit intersubunit contact surface. Of these sequences, 60 Denatured-only and six Overlap peptides had resolved coordinates in the structure. A mapped peptide was defined as a Factor B interface peptide if at least one of its heavy atoms lay within 4.5 Å of a Factor B heavy atom. This criterion identified 13 Denatured-only interface peptides and no Overlap interface peptides. The sequence map placed the interface peptides in multiple segments across the C3b β and α′ chains (**Figure 3a**). The whole-complex views located these segments along the Factor B contact surface and the mapped Overlap peptides outside this interface, while the five local views illustrated the residue-level contacts with Factor B (**Figure 3b, 3c**). Burial analysis quantified the shielding associated with these contacts. The 13 Denatured-only interface peptides had a median Factor B-induced burial fraction of 16%, compared with median values of 0% for the remaining 47 mapped Denatured-only peptides and the six mapped Overlap peptides (**Supplementary Table S3**). Thus, the Factor B interface peptides belonged exclusively to the Denatured-only class, and their positions corresponded to regions with greater burial upon association with Factor B. Together, the mapping and burial results connect a specific subset of the experimentally observed detection differences to local shielding at a partner-binding surface. Complete peptide mapping results, domain annotations^22^, and structural measurements are provided in **Supplementary Table S3**.

**Figure 3.**
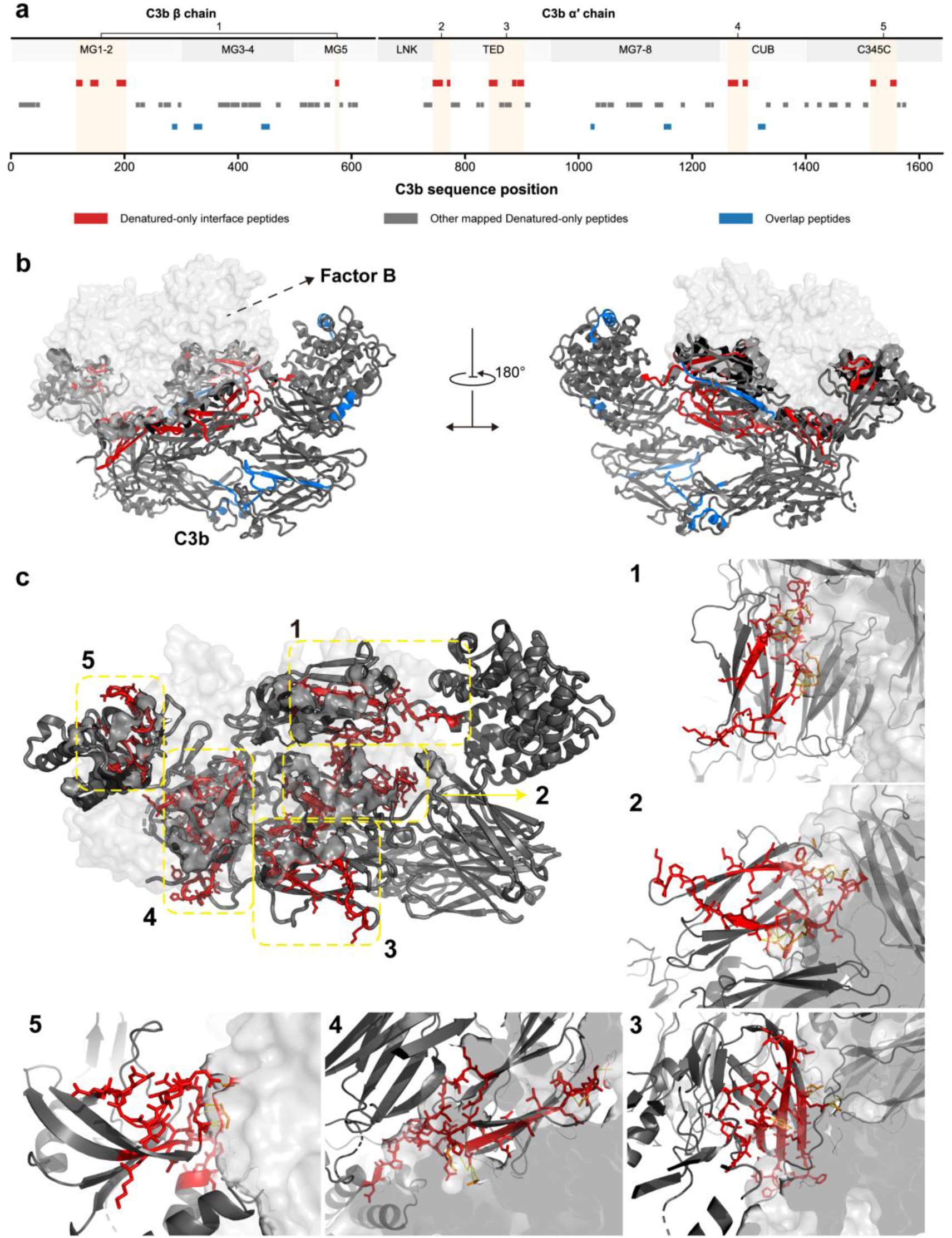
Mapping of C3b peptides onto the C3b–Factor B complex (PDB 2XWJ). (a) Peptide positions along the C3b β and α′ chains, with domain annotations. Red marks Denatured-only interface peptides, gray marks other mapped Denatured-only peptides, and blue marks Overlap peptides. Numbered regions 1–5 correspond to the local views in panel c. (b) Two views, rotated by 180°, of the C3b–Factor B complex. C3b is shown as dark ribbons and Factor B as a pale surface; mapped peptide regions are colored by detection class. (c) The overall view shows the five interface regions (yellow dashed boxes) and the corresponding close-up views. Red peptides denote Denatured-only interface peptides, and blue peptides denote Overlap peptides. In the close-up views, yellow dashed lines indicate distances of <3.5 Å between heavy atoms, and orange-colored residues highlight the interface residues. All 13 interface peptides identified using the ≤4.5 Å heavy-atom criterion were Denatured-only.

The identified interface peptides include regions implicated in the activation of the complement proconvertase. A subset overlaps the macroglobulin 2 (MG2) and CUB regions of C3b, linking the observed Denatured-only pattern to a functionally characterized part of the Factor B interface (**Supplementary Table S3**). Forneris et al. showed that the serine protease domain of Factor B contacts these domains in the open C3bB proconvertase and that mutations at the Factor B contact site reduce Factor D-mediated cleavage and convertase formation.^19^This cleavage generates the active alternative-pathway C3 convertase, C3bBb^23,24^.The overlap between the mapped peptides and these regions therefore places part of the Denatured-only peptide set at a contact surface involved in this activation process. This case demonstrates that PLAPI can connect peptide detection differences to both local shielding by a protein partner and an assembly surface with an established role in complement activation.

### A2M Intersubunit Contacts and Bait-Region Accessibility

A2M was selected as a complementary structural case because it contains a protease-sensitive bait region with an established role in protease-triggered activation^25^, and the A2M dataset included both Denatured-only and Overlap peptides. The dataset contained 56 Denatured-only and 3 Overlap peptide sequences; in PDB 6TAV^25^, 55 and 1 sequences, respectively, had resolved coordinates in at least one chain copy (**Supplementary Table S4**). For intersubunit mapping, peptides were mapped onto chain A and assessed against neighboring chains B, C, and D using the same interface-peptide criterion described above. This identified 12 Denatured-only interface-peptide records corresponding to 11 distinct contact regions, with two records mapping to the same region. Eight regions contacted chain C, two contacted chain B, and one contacted chain D (**Figure 4a**; **Supplementary Table S4**). **Figure 4a** locates the regions along the chain A sequence, **Figure 4b** shows their distribution across the tetramer, and **Figure 4c** provides close-up views of the A–B, A–C, and A–D contacts. Thus, the identified Denatured-only interface peptides map to multiple intersubunit surfaces in the A2M tetramer. Complete peptide sequences, detection classes, coordinate coverage, contacting-chain assignments, domain annotations, and local-view groupings are provided in **Supplementary Table S4**.

**Figure 4.**
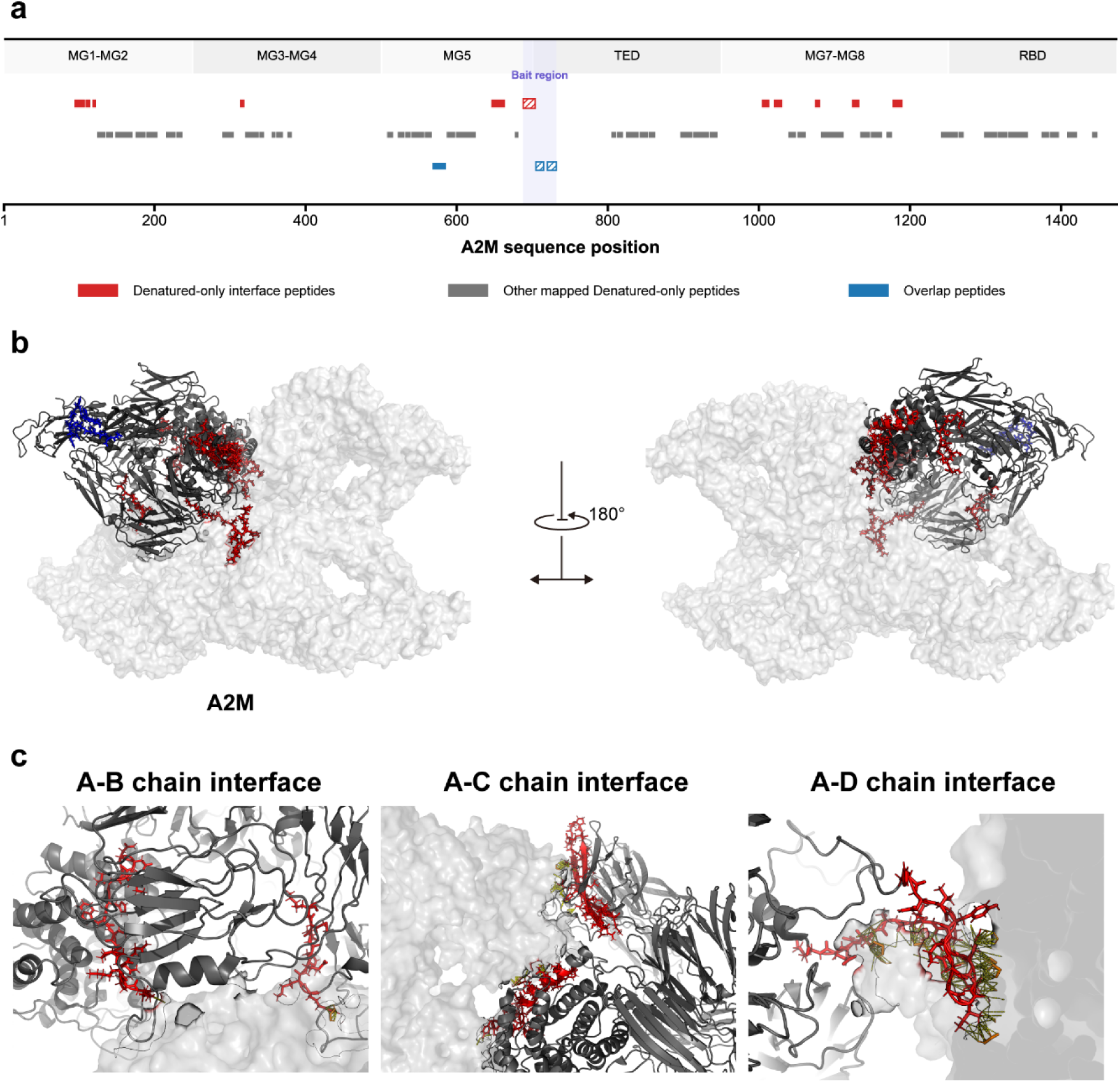
A2M peptide mapping, intersubunit contacts, and bait-region detection patterns (PDB 6TAV). (a) Peptide positions along the A2M sequence with domain annotations. Red marks Denatured-only interface peptides, gray marks other mapped Denatured-only peptides, and blue marks Overlap peptides. The shaded band indicates the bait region; hatched portions denote sequence lacking resolved coordinates in the reference structure. (b) Two views of the A2M tetramer, rotated by 180°, showing mapped peptide regions on chain A in the context of the other subunits. (c) Close-up views of the A–B, A–C, and A–D interfaces. Yellow dashed lines indicate distances of <3.5 Å between heavy atoms, and orange-colored residues highlight the interface residues. Twelve Denatured-only interface-peptide records correspond to 11 distinct contact regions. In the bait region, the Denatured-only peptide at residues 688–704 is adjacent to Overlap peptides at residues 705–715 and 720–732.

The bait-region results reveal different detection states among adjacent peptides in a functionally characterized protease-triggering element. Two of the three A2M Overlap peptides mapped to residues 705–715 and 720–732, while the neighboring 688–704 peptide was Denatured-only (**Figure 4a**; **Supplementary Table S4**). The resolved portion of 688–704 includes residues at an A–C contact (**Figure 4c**). Previous studies showed that cleavage of the bait region initiates the conformational transition that traps the attacking protease^25,26^. In this functional context, the comparison shows that Native digestion recovered two bait-region segments, whereas the adjacent 688–704 segment was detected reproducibly only under Denatured conditions. It therefore resolves peptide-level differences in recovery across neighboring segments of a known protease-triggering region.

### STRING Association Network and Functional Enrichment

STRING analysis^27,28^ identified high-confidence associations among proteins carrying Denatured-only peptides. The 129 candidate proteins supported by at least one Denatured-only peptide were submitted to STRING, which mapped 107 of them. Applying a combined-score cutoff of ≥0.90 yielded the network shown in **Figure 5a**, comprising 76 nodes and 167 edges with scores ranging from 0.900 to 0.999.These associations involved complement components, apolipoproteins, coagulation factors, and serine-protease regulators, organizing the connected proteins into groups with established roles in immune defense, lipid transport, and hemostasis. Thus, proteins selected through peptide-level classification were linked by known functional associations across several serum biological processes. The complete input list and supporting peptide evidence are provided in **Supplementary Table S5**; STRING protein pairs with combined scores ≥0.90 are listed in **Supplementary Table S6.**

**Figure 5.**
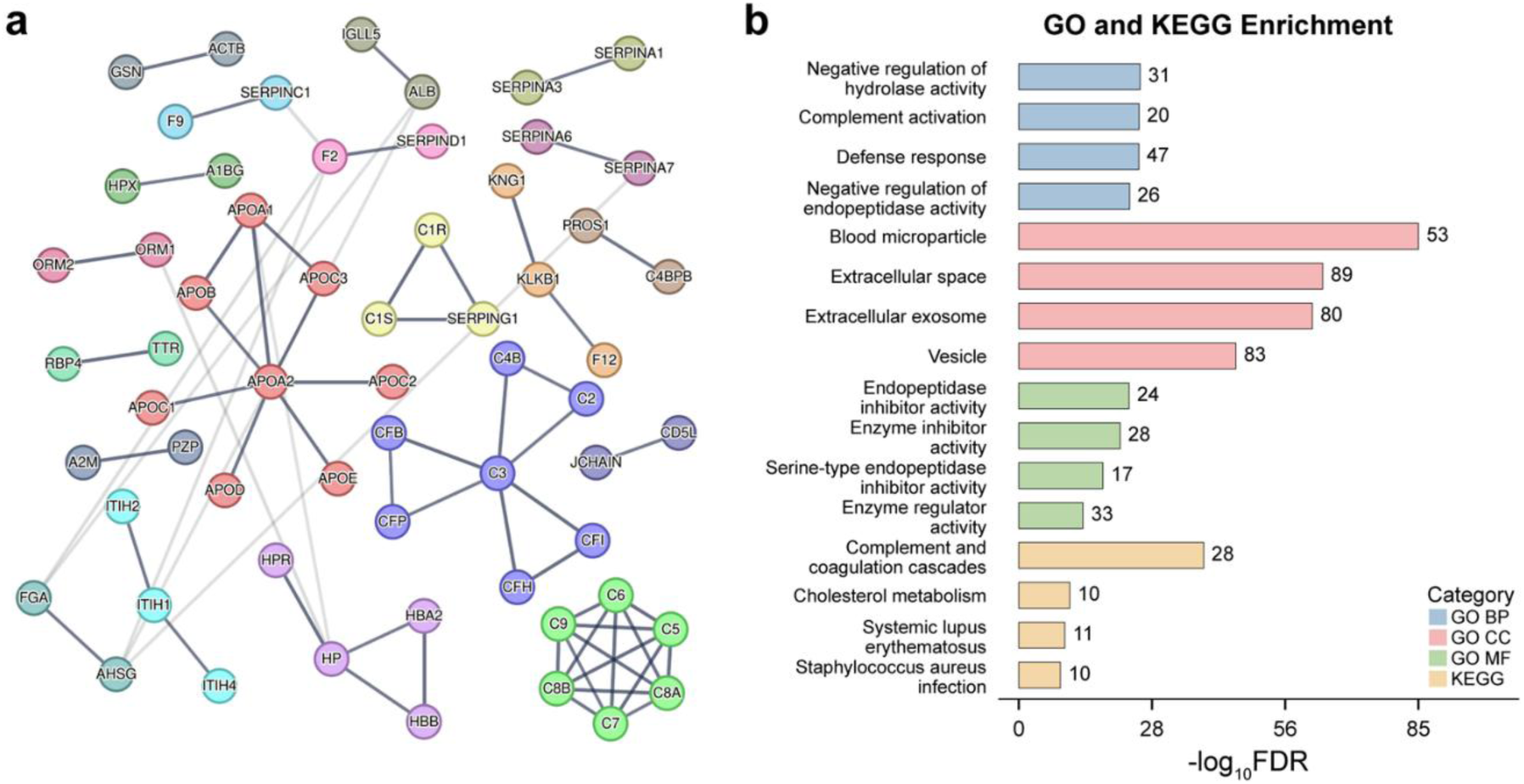
STRING associations and functional enrichment among proteins carrying Denatured-only peptides. (a) The network of 76 connected proteins and 167 associations retained at a STRING combined-score threshold of ≥0.90. Nodes represent proteins and edges represent STRING associations. (b) The four displayed terms in each GO Biological Process, Cellular Component, and Molecular Function category and the four displayed KEGG pathways from enrichment analysis of the 107 proteins mapped to STRING. Bar length represents −log₁₀(FDR); numbers beside bars indicate the number of input proteins associated with each term or pathway. Colors identify the enrichment category as indicated in the legend.

Functional enrichment of the same 107-protein input set highlighted extracellular defense and protease regulation (**Figure 5b**). Gene Ontology (GO)^29^ biological-process terms included complement activation, defense response, and negative regulation of hydrolase activity, involving 20, 47, and 31 proteins, respectively. In the molecular-function category, endopeptidase inhibitor activity and serine-type endopeptidase inhibitor activity involved 24 and 17 proteins, respectively. Together, these annotations link the protein set to both immune defense and the regulation of proteolytic activity. Cellular-component enrichment further placed these functions in an extracellular context, with 89 proteins annotated to extracellular space and 53 to blood microparticle. At the pathway level, Kyoto Encyclopedia of Genes and Genomes (KEGG)^30^ analysis identified complement and coagulation cascades as the most significantly enriched pathway, involving 28 proteins with a false discovery rate (FDR) of 6.92 × 10⁻⁴⁰. Cholesterol metabolism was also enriched, involving 10 proteins with an FDR of 1.66 × 10⁻¹¹. These pathway results connect the complement, coagulation, and apolipoprotein groups observed in the network to their associated biological processes. Collectively, the enrichment results identify extracellular defense and protease regulation as major functional themes among proteins carrying Denatured-only peptides, alongside lipid metabolism. Complete GO and KEGG enrichment results are provided in **Supplementary Table S7**.

The network and enrichment results place the C3b and A2M structural observations within the broader functional context of serum protease regulation. In C3b, Denatured-only interface peptides mapped to a Factor B contact surface involved in complement convertase assembly. In A2M, adjacent peptides with different detection states occurred within the bait region that initiates protease capture. These observations concern two processes represented in the enrichment results: complement activation and protease inhibition. The C3b and A2M cases therefore provide peptide-level observations within functional regions of proteins involved in these broader processes. Together, the structural and functional analyses relate the observed peptide detection patterns to regions involved in extracellular defense and the control of protease activity.

## METHODOLOGICAL IMPLICATIONS AND SCOPE

PLAPI uses differences in peptide profiles to highlight regions of endogenous serum proteins with condition-dependent peptide recovery. The Native arm captures peptide release during brief digestion under native conditions, while the Denatured arm provides a reference profile after unfolding and extended digestion. Profile differences reflect local protease accessibility and digestion kinetics; peptide recovery and MS detectability also shape the observed signal. Mapping the peptides to reference structures places these patterns within assembled proteins.

The workflow’s main methodological advantage is its capacity to survey candidate regions in complex endogenous samples before a specific interaction is selected. Relative to AP-MS, it adds peptide-level regional evidence of condition-dependent recovery to protein-level interaction evidence^1,6^. FLiP-MS separates assembly states to derive peptide markers, whereas Holfeld et al. measure responses to selected purified bait proteins^15,16^. The present workflow complements these approaches by enabling regional discovery without either prerequisite.

This regional readout helps prioritize segments within candidate proteins for follow-up. The resulting assignments can guide the selection of proteins and peptides for targeted measurements and orthogonal PPI studies, moving the analysis from proteome-wide observations to focused examination of candidate interfaces and functional regions.

The current analysis was based on one serum sample and three technical preparation replicates, and structural interpretation relied on published reference assemblies. Analyses of independent serum samples and controlled changes in protein assembly or activation can help assess the reproducibility and scope of this readout.

## CONCLUSION

Taken together, PLAPI provides a peptide-level strategy for prioritizing candidate structural regions in complex serum proteomes. Its value lies in connecting condition-dependent peptide recovery with structural context, thereby guiding focused investigation of potential protein interfaces and functional regions. Further validation using independent serum specimens and orthogonal approaches will be important to establish the biological relevance of the identified regions.

---

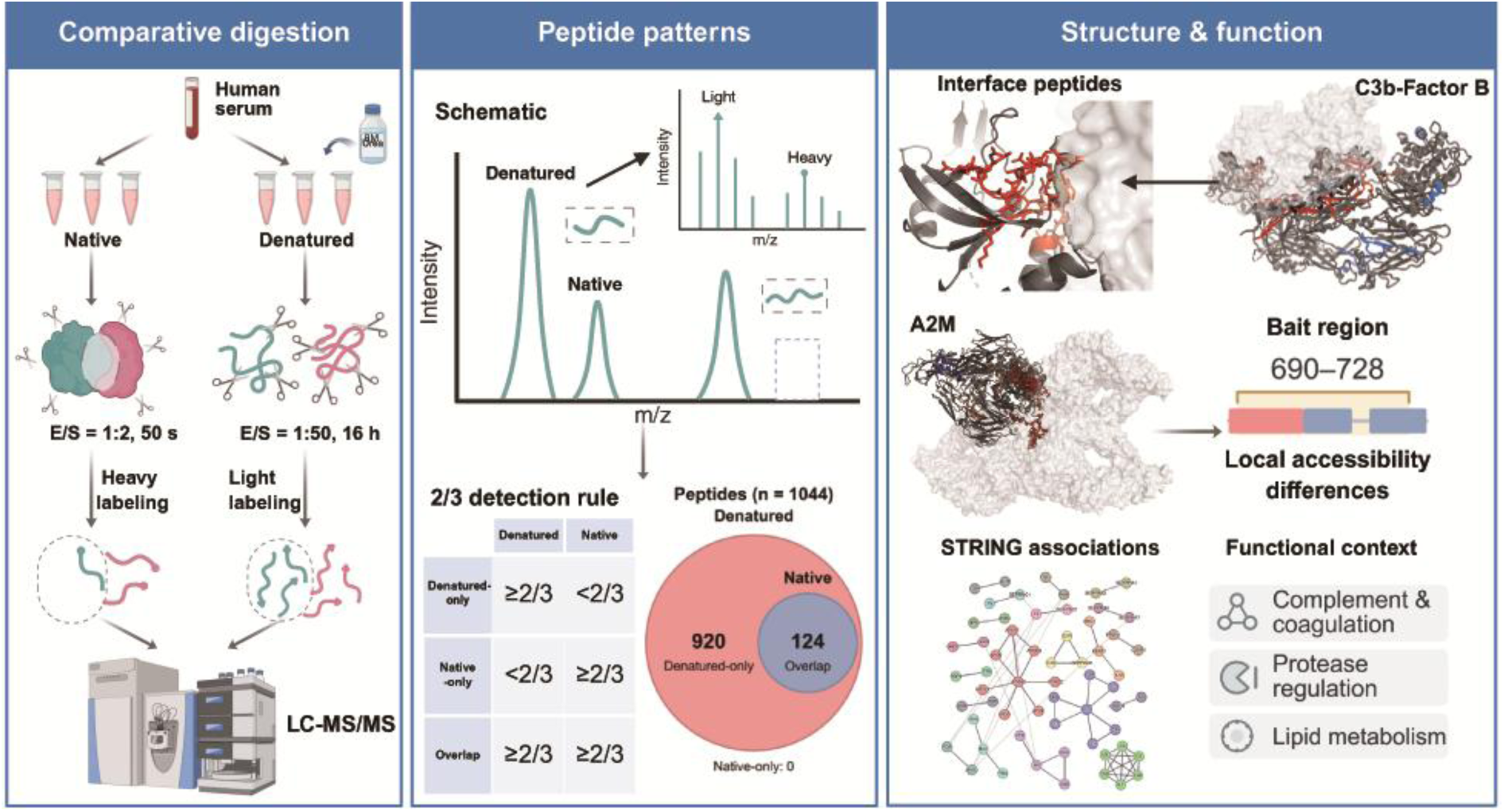
TOC Graphic.

