## Supplementary material for "Peptide-level assay of protein interaction using combinatorial native and denatured proteolysis": Supplemental information_PLAPI.docx

### S1. Supplementary Experimental Methods

#### S1.1. Reduction and alkylation

For reduction, DTT was added to a final concentration of 10 mM, followed by incubation at 55 °C for 30 min. After cooling to room temperature, IAA was added to a final concentration of 20 mM, and samples were incubated in the dark for 30 min. A second DTT addition, equal to the first, was followed by incubation at room temperature for 30 min. Native samples underwent these steps after GdmCl quenching; Denatured samples underwent them before trypsin digestion.

#### S1.2. Denatured digestion

For each Denatured preparation, 2 μL of serum was mixed with 8 μL of 8 M urea, vortexed for 10 s, and incubated at room temperature for 1 h. The mixture was diluted to 100 μL with 90 μL of PBS. DTT and IAA were added as described in Section S1.1. Trypsin was added at an enzyme-to-substrate mass ratio (E:S) of 1:50, and samples were incubated at 37 °C for 16 h. Digestion was terminated by adding TFA to a final concentration of 0.1%, followed by vortex mixing for 10 s. Peptides were collected using the same ultrafiltration procedure as for Native samples (Section S1.3).

#### S1.3. Ultrafiltration and peptide collection

Both sample types were transferred to 10 kDa centrifugal filter units and centrifuged at 14,000 × g for 1 h. The filter membrane was washed with 200 μL of 60 mM HEPES, pH 8.2, and centrifuged again at 14,000 × g for 1 h. The filtrates from both centrifugation steps were combined and acidified by adding TFA at 1% of the filtrate volume before C18 desalting.

#### S1.4. C18 desalting and stable isotopic diethyl labeling

Peptide mixtures were desalted using Jupiter C18 material (15 μm, 300 Å; Phenomenex). Before labeling, the C18 material was activated three times with 100 μL of acetonitrile and equilibrated three times with 100 μL of 0.1% TFA. After sample loading, it was washed eight times with 100 μL of 0.1% TFA. Peptides were eluted four times with 100 μL of 50% acetonitrile/0.1% TFA and four times with 100 μL of 80% acetonitrile/0.1% TFA. Eluates were combined and dried by vacuum centrifugation.

Dried peptides were reconstituted in 100 μL of trifluoroethanol. For reductive diethylation, 20% light acetaldehyde (CH₃CHO) or heavy [U-¹³C₂]acetaldehyde (¹³CH₃¹³CHO) was added at 0.25 μL per 1 μg of peptide. Samples were mixed for 1 min, an equal volume of freshly prepared 0.6 M sodium cyanoborohydride was added, and samples were mixed for another minute. Labeling proceeded at 37 °C for 1 h. The reaction was quenched with 4% ammonium hydroxide at a volume equal to that of the acetaldehyde labeling reagent added to that reaction. Corresponding light-labeled Denatured and heavy-labeled Native preparations were directly combined. The combined sample was desalted using C18 material, with 50 μL solvent portions for activation, equilibration, washing, and elution, and then dried by vacuum centrifugation.^1^

The net mass additions per modified amino group were +56.0626 Da for light diethylation (C₄H₈) and +60.0760 Da for heavy diethylation (¹³C₄H₈), a heavy–light difference of 4.0134 Da. These modifications were assigned to peptide N termini and lysine residues.

#### S1.5. LC–DDA–MS/MS acquisition

Peptides (5 μg per analysis) were analyzed using a nano-ESI Q Exactive mass spectrometer coupled to a Dionex Ultimate 3000 RSLCnano system. Samples were loaded onto a self-packed C18 trap column (200 μm i.d. × 5 cm, 360 μm o.d.) packed with Phenomenex Jupiter C18 particles (5 μm, 300 Å) at 5 μL/min for 5 min. Peptide separation was performed using a self-packed C18 analytical column (75 μm i.d. × 70 cm, 360 μm o.d.) packed with the same C18 material at a flow rate of 300 nL/min. Mobile phase A consisted of 99.8% water and 0.2% formic acid, whereas mobile phase B consisted of 90% acetonitrile, 9.8% water, and 0.2% formic acid. The gradient and re-equilibration program was as follows: 7–42% B over 95 min, 42–95% B over 5 min, 95% B for 3 min, 95–7% B over 2 min, and 7% B for 10 min. The 115-min gradient and re-equilibration sequence, preceded by the 5-min loading step, constituted a total LC program of 120 min.

The electrospray voltage was 2.8 kV, and the ion-transfer tube temperature was 300 °C. MS1 scans were acquired over an m/z range of 700–2,000 at a resolution of 60,000, with an automatic gain control (AGC) target of 3.0 × 10⁶ and a maximum injection time of 50 ms. Precursor ions with charge states of 1+ and greater than 7+ were excluded from selection. The top 20 most abundant precursor ions were selected for HCD-MS/MS analysis (Top20 data-dependent acquisition) at a resolution of 30,000, with an AGC target of 5.0 × 10⁵, an isolation window of 3.0 m/z, and a maximum injection time of 250 ms. Stepped normalized collision energies (NCEs) of 20%, 30%, and 40% were applied, and dynamic exclusion was set to 30 s.

#### S1.6. Database searching and peptide quantification

Raw files were processed using Proteome Discoverer version 1.4.0.288 against a local human plasma protein FASTA database (uniprot plasma-6274_20201105.fasta). Two separate Sequest HT search nodes were used for the light- and heavy-labeled peptide datasets. Trypsin (full specificity) was selected as the digestion enzyme, allowing up to two missed cleavages. The minimum and maximum peptide lengths were set to 6 and 144 amino acids, respectively, with a maximum ΔCn of 0.05. Precursor and fragment mass tolerances were set to 10 ppm and 0.6 Da, respectively.

For light-labeled peptides, fixed modifications included diethylation (+56.063 Da) at peptide N termini and lysine residues and carbamidomethylation (+57.021 Da) at cysteine residues. For heavy-labeled peptides, the corresponding fixed diethylation modification was +60.076 Da at peptide N termini and lysine residues, with carbamidomethylation (+57.021 Da) at cysteine residues. No dynamic modifications were specified. The maximum number of equal modifications per peptide was set to three. Peptide-spectrum matches were validated using the Percolator node, with both strict and relaxed target FDR thresholds set to 0.01 and validation based on q-values.

Precursor-ion quantification was performed using the Precursor Ion Quantifier node. Paired light- and heavy-labeled precursor intensities were used to calculate within-preparation Denatured/Native ratios, defined as log₂(Denatured/Native). Missing quantitative values were not imputed for replicate-based peptide detection classification, as described in Section S2.1.

### S2. Data Analysis and Calculation Definitions

#### S2.1. Replicate-based peptide classification

Peptide detection was assessed separately across three technical preparations per condition from the same serum sample. Within an individual preparation, a peptide was considered detected when at least one corresponding peptide-spectrum match passed Percolator validation at the 1% FDR threshold based on q-values. A peptide was considered reproducibly detected in a condition when it was detected in at least two of the three preparations.

Denatured-only peptides met this threshold in Denatured preparations but not in Native preparations. Overlap peptides met the threshold in both conditions. Native-only peptides met the threshold only in Native preparations. Peptides meeting the threshold in neither condition were excluded from the reproducibly detected set and are labeled Unreliable in Table S1. Thus, Denatured-only permits one Native detection; it does not require absence from all three Native preparations.

For paired quantitative measurements, log₂FC was calculated as log₂(Denatured/Native). Missing quantitative values were not imputed for detection classification. Replicate-level detections and category assignments are provided in Table S1.

#### S2.2. Sequence features

For sequence features, let $n$ denote peptide length. K/R density was calculated as $10N_{\mathrm{KR}}/n$ (K/R residues per 10 amino acids); acidic-residue fraction as $N_{\mathrm{DE}}/n$; hydrophobic-residue fraction as $N_{\mathrm{AVILMFWYP}}/n$; and the net charge proxy as $N_{\mathrm{KR}}-N_{\mathrm{DE}}$. The acidic-neighbor fraction was calculated as the fraction of K/R sites with at least one D/E residue within two sequence positions on either side. The proline-neighbor score was calculated as the number of immediately adjacent proline residues per K/R site, counting the two sides separately. A proline adjacent to two K/R sites was counted for each site. Both neighbor metrics were set to zero for peptides without K/R residues.

#### S2.3. Reference sequences and structural mapping

Peptides were mapped to UniProt reference sequences. For the focused structural analyses, C3-derived peptides (P01024) were mapped to the C3b–Factor B reference structure (PDB 2XWJ), and A2M-derived peptides (P01023) were mapped to the tetramer in PDB 6TAV. Peptide positions, coordinate coverage, and chain assignments are reported in Tables S3 and S4.^2–4^

Atom-based structural metrics used the residues with resolved coordinates. Global feature comparisons used the available peptide–protein records in Table S2; records without an unambiguous value for a metric were omitted from that comparison. Focused interface mapping and global structural-feature comparisons were treated as separate analyses.

#### S2.4. Relative solvent-accessible surface area

For residue $i$, relative solvent-accessible surface area was calculated as $\mathrm{rSAS}A_{i}=SASA_{i}/SASA_{max,i}$, using residue-specific maximum accessible areas: A 129, R 274, N 195, D 193, C 167, Q 225, E 223, G 104, H 224, I 197, L 201, K 236, M 224, F 240, P 159, S 155, T 172, W 285, Y 263, and V 174 Å². Peptide mean rSASA was the arithmetic mean across mapped residues. A residue was classified as exposed at rSASA ≥0.25; the exposed fraction was the number of exposed mapped residues divided by the number of mapped residues. SASA was calculated in PyMOL using a solvent radius of 1.4 Å and dot density 4. The PyMOL version was 3.1.^5^

#### S2.5. Global relative Cα B-factor

For the global B-factor comparison in Table S2, the mean Cα B-factor of each mapped peptide was divided by the mean Cα B-factor across all chains of its reference structure. Only Cα atoms with coordinates in the first structural model were included. Normalized values were then averaged across valid mappings of each peptide–protein entry:

$B_{rel,p,k}=\frac{\overline{B}_{p,k}}{\overline{B}_{s(k)}}, B_{rel,p}=\frac{1}{K_{p}}\sum_{k=1}^{K_{p}} B_{rel,p,k}$	(S1)

Here, $\overline{B}_{p,k}$ is the mean peptide Cα B-factor for mapping k, $\overline{B}_{s(k)}$ is the mean Cα B-factor of the corresponding reference structure, and $K_{p}$ is the number of valid mappings.

#### S2.6. Interface contacts and C3b-specific structural metrics

An interface peptide was defined by a minimum distance of ≤4.5 Å between any peptide non-hydrogen atom and any non-hydrogen atom in the relevant neighboring chain. Factor B was the partner for the C3b analysis. For A2M, peptides mapped to chain A were assessed for contacts with neighboring chains B, C, and D. Interface assignments are listed in Tables S3 and S4.

For C3b structure metrics, relative Cα B-factor was calculated as the mean peptide Cα B-factor divided by the mean Cα B-factor of the corresponding C3b chain. For each mapped chain copy, Factor B-induced burial fraction was calculated as

$f_{buried,p,k}=\frac{A_{iso,p,k}-A_{complex,p,k}}{A_{iso,p,k}}$	(S2)

In eq S2, $A_{iso,p,k}$ and $A_{complex,p,k}$ are the SASA values of peptide p in mapped copy k of the isolated C3b chain and the C3b–Factor B complex, respectively. The burial fraction and other peptide metrics were calculated for each mapped copy and then averaged across copies for each peptide. The chain-normalized C3b B-factor in Table S3 is distinct from the whole-structure-normalized global metric in Table S2.

#### S2.7. Global feature comparisons

Global sequence and structural feature distributions were compared using two-sided Mann–Whitney U tests with tie and continuity corrections. Each test used the available values for that metric; missing values were excluded without imputation. Unadjusted P values are reported. Sample sizes denote peptide records, not biological replicates. The comparisons are shown in Figure S1, with underlying values in Table S2.

#### S2.8. STRING associations and functional enrichment

Proteins assigned at least one Denatured-only peptide were analyzed using the STRING website for Homo sapiens. STRING analysis was performed using STRING version 12.5 for Homo sapiens (accessed on November 1, 2026)). Of 129 canonical UniProt candidate identifiers, 107 were mapped to STRING identifiers (Table S5). Protein assignments supported by shared peptides are retained in the candidate-protein table.^6^

The network shown in main-text Figure 5a used a combined-score threshold of ≥0.90 and contained 76 connected nodes and 167 edges. Table S6 lists the retained associations. GO Biological Process, Molecular Function, and Cellular Component terms and KEGG pathways were obtained from the STRING enrichment output for the 107 mapped proteins, using the entire genome proteome. STRING-reported FDR values and annotated-protein counts were retained, and the top four terms in each displayed category were ranked by FDR. Table S7 includes the full returned GO and KEGG results; no additional edge-confidence filter was applied to that table.

#### S2.9. Global feature comparison

No significant distributional differences were observed between Denatured-only and Overlap peptides for the sequence features, mean rSASA, exposed-residue fraction, or relative Cα B-factor in the available records (all unadjusted P > 0.05; Figure S1 and Table S2). These global comparisons did not identify a consistent distinction between the detection classes. The focused structural analyses therefore examined peptide locations within individual protein assemblies.

### S3. Supplementary Figure

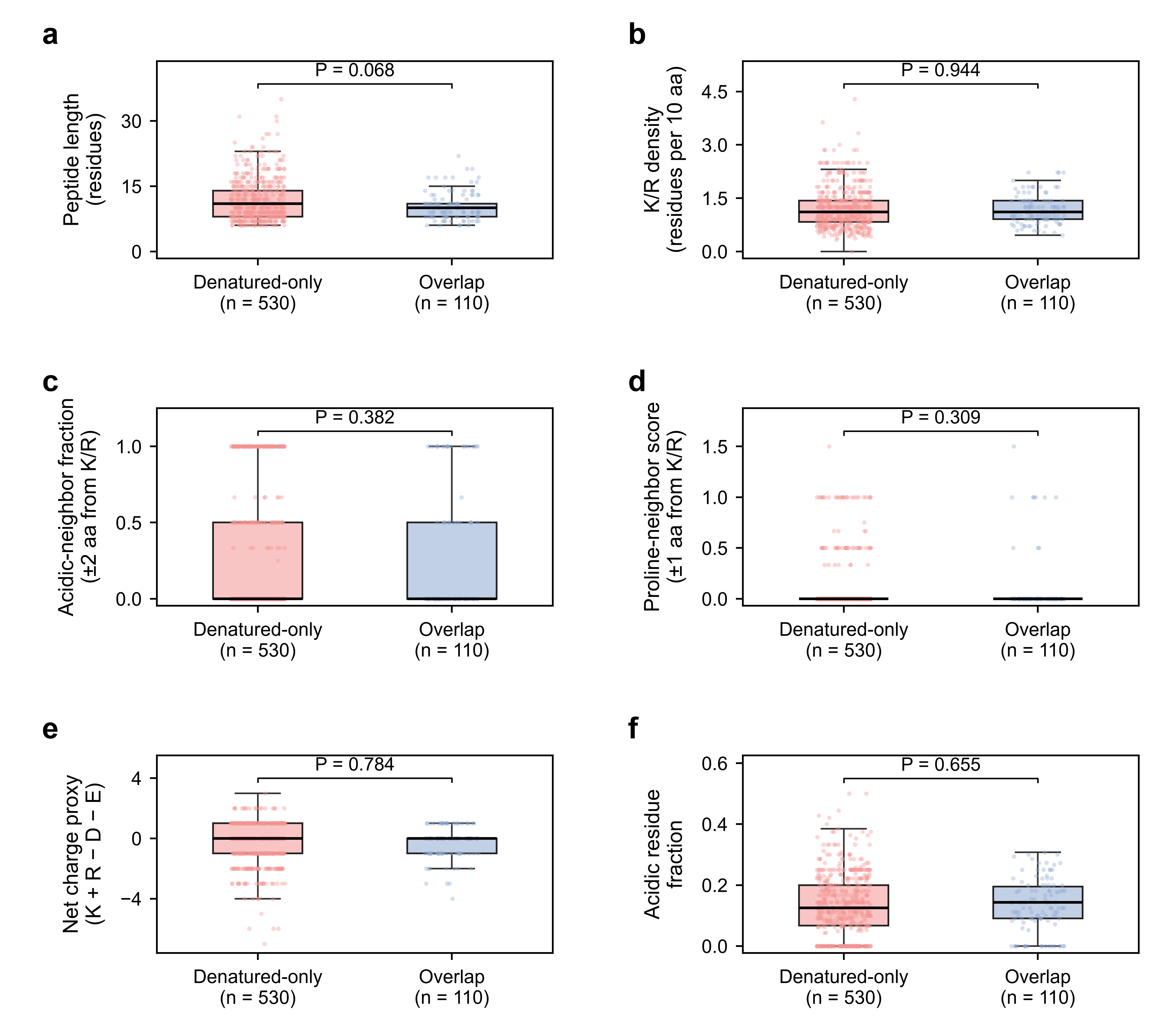

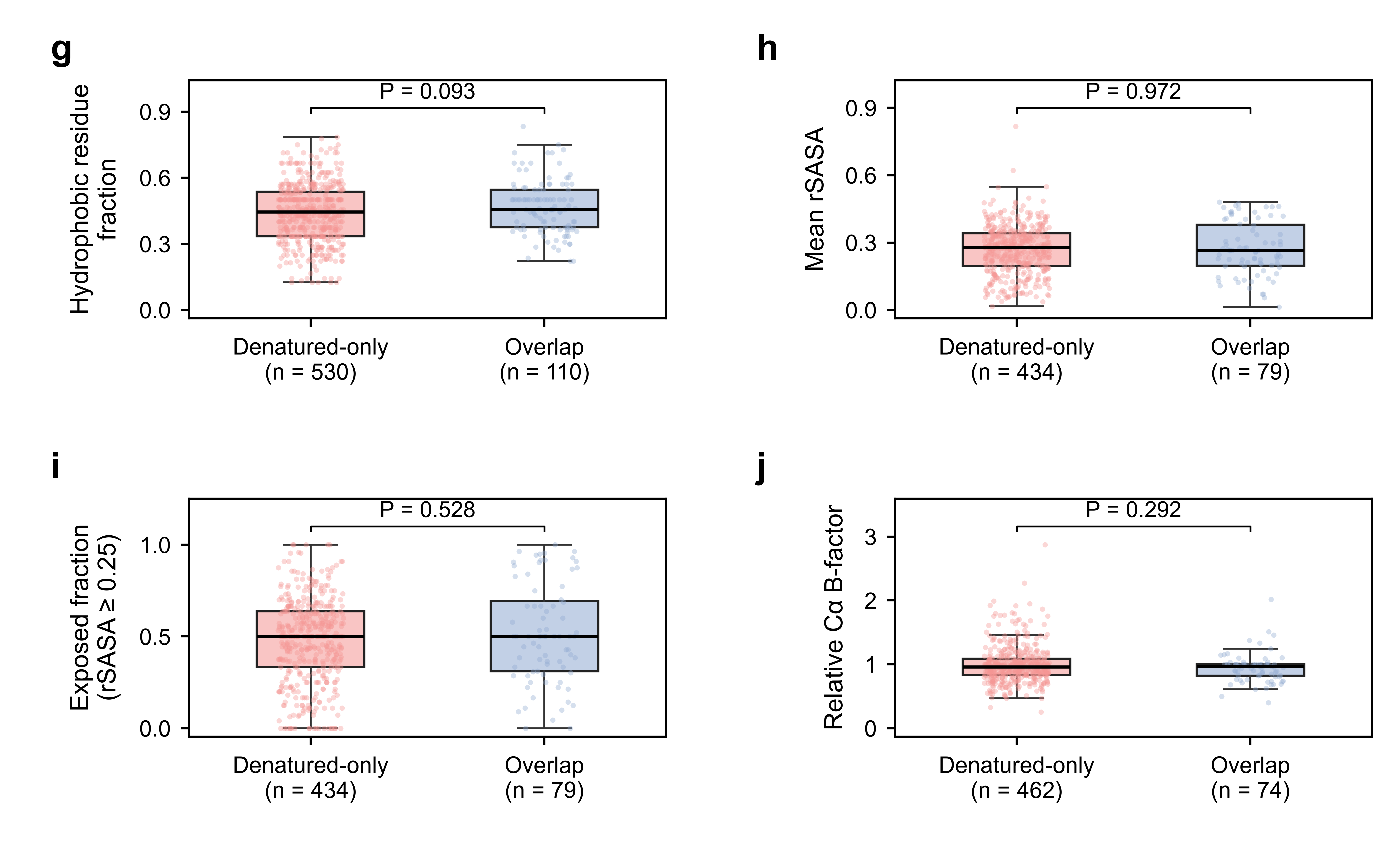

**Figure S**1**. Global sequence and structural features of Denatured-only and Overlap peptides. Panels a–f.**

(a) Peptide length. (b) K/R density, expressed as K/R residues per 10 amino acids. (c) Acidic-neighbor fraction: the fraction of K/R sites with at least one D/E residue within two sequence positions on either side. (d) Proline-neighbor score: immediately adjacent proline residues per K/R site, counting both sides separately. Neighbor metrics were set to zero for peptides without K/R residues. (e) Net charge proxy: the number of K/R residues minus the number of D/E residues. (f) Acidic-residue fraction. Coral, Denatured-only; blue, Overlap. Points represent individual peptide records. Boxes show the interquartile range (IQR), with medians as center lines. Whiskers extend to the most extreme observations within 1.5 × IQR; all observations, including those beyond the whiskers, are shown. In panel d, both groups have a first quartile, median, and third quartile of zero, so the boxes collapse to a line. P values are unadjusted, two-sided Mann–Whitney U test results. Data: Table S2. Panels g–j continue on the next page. (g) Hydrophobic-residue fraction, using A, V, I, L, M, F, W, Y, and P. (h) Mean relative solvent-accessible surface area (rSASA). (i) Fraction of mapped residues with rSASA ≥ 0.25. (j) Relative Cα B-factor: the peptide mean divided by the mean across all coordinate-bearing Cα atoms in the first model of the corresponding reference structure, averaged across valid mappings of each peptide–protein entry. Colors and box-plot conventions are the same as in panels a–f. Sample sizes (n) indicate the available peptide records for each metric. P values are from two-sided Mann–Whitney U tests and are unadjusted for multiple comparisons. Underlying data are provided in Table S2.

### S4. Supplementary Tables

Tables S1–S7 are supplied as separate worksheets in the accompanying spreadsheet file. Each worksheet contains its title, column headings, and explanatory notes.

**Table S1. Replicate-level peptide detections and category assignments.** Presence/absence calls for each Native and Denatured preparation, protein assignments, and categories under the 2-of-3 rule. The table retains the Unreliable records so that the classification can be traced.

**Table S2. Sequence and structural features.** Sequence features, rSASA metrics, and the independent global relative Cα B-factor distribution used for Figure S1. Metric-specific availability and mapping sources are retained. Global B-factor normalization uses the reference structure across all chains.

**Table S3. C3-derived peptide mapping onto C3b–Factor B.** Peptide sequences, detection classes, coordinate coverage, interface assignments, and structural metrics in PDB 2XWJ, including Factor B-induced burial fractions and C3b chain-normalized B-factors.

**Table S4. A2M peptide mapping and intersubunit contacts.** Peptide sequences, detection classes, coordinate coverage, contacting chains, domain annotations, and local-view assignments for PDB 6TAV.

**Table S5. Candidate proteins and STRING identifier reconciliation.** The 129 canonical UniProt candidates supported by Denatured-only peptides, their peptide evidence, and mapping to 107 STRING identifiers. Unmapped identifiers and mapped nodes without edges are distinguished.

**Table S6. STRING associations retained for Figure 5a.** The 167 protein associations with combined scores ≥0.90 that form the 76-node network in main-text Figure 5a.

**Table S7. GO and KEGG enrichment results.** The returned GO Biological Process, Molecular Function, Cellular Component, and KEGG terms for the 107 mapped proteins, including STRING-reported FDR values, term-associated protein counts, and matched proteins.
